# Causal Brain Network Evaluates Surgical Outcomes in Patients with Drug-Resistant Epilepsy

**DOI:** 10.1101/2024.03.03.583165

**Authors:** Yalin Wang, Minghui Liu, Wentao Lin, Weihao Zheng, Tiancheng Wang, Yaqing Liu, Hong Peng, Wei Chen, Bin Hu

## Abstract

Network neuroscience has greatly facilitated epilepsy studies; meanwhile, drug-resistant epilepsy (DRE) is increasingly recognized as a brain network disorder. Unfortunately, surgical success rates in patients with DRE are still very limited, varying 30% ∼ 70%. At present, there is almost no systematic exploration of intracranial electrophysiological brain network closely related to surgical outcomes, and it is not clear which brain network methodologies can effectively promote DRE precision medicine. In this retrospective comparative study, we included multicenter datasets, containing electrocorticogram (ECoG) data from 17 DRE patients with 55 seizures. Ictal ECoG within clinically-annotated epileptogenic zone (EZ) and non-epileptogenic zone (NEZ) were separately computed using six different algorithms to construct causal brain networks. All the brain network results were divided into two groups, successful and failed surgery. Statistical results based on the Mann-Whitney-U-test show that: causal connectivity of α-frequency band (8 ∼ 13 Hz) in EZ calculated by convergent cross mapping (CCM) gains the most significant differences between the surgical success and failure groups, with a *P* value of 7.85e-08 and Cohen’s d effect size of 0.77. CCM-defined EZ brain network can also distinguish the successful and failed surgeries considering clinical covariates (clinical centers, DRE types) with *P* < 0. 001. Based on the brain network features, machine learning models are established to predict the surgical outcomes. Among them, SVM classifier with Gaussian kernel function and Bayesian Optimization demonstrates the highest average accuracy of 84.48% through 5-fold cross validation, further indicating that the CCM-defined EZ brain network is a reliable biomarker for predicting DRE’s surgical outcomes.

## I Introduction

### A. Background

Epilepsy is one of the most common neurological disorders, affecting over 60 million individuals worldwide [1], [2]. There is approximately one-third of epilepsy patients who are resistant to antiepileptic medications [3], [4], [5], called “drug resistant epilepsy (DRE)”. The DRE patients are defined as continued seizures despite two trials of appropriately chosen anti-epileptic drugs [6], [7], [8]. For patients with DRE, neurosurgery is the most effective treatment. Successful surgery requires complete removal or disconnection of the epileptogenic zone (EZ), while current surgical success rates are very low. Only approximately 30 ∼ 70% of the DRE patients undergo surgery achieve seizure-freedom [9]. Surgical failure can lead to potential consequences such as permanent nerve damage, and persistent seizures in children will lead to neurodevelopmental dysplasia. Expensive surgical cost is also a heavy financial burden. Therefore, exploring biomarkers for surgical outcomes and predicting surgical outcomes before neurosurgery has attracted considerable attention among clinicians [10], [11], [12], [13].

Epilepsy is growingly considered as a network-related disorder that cannot be entirely defined by the isolated analysis of neurophysiological signals [9]. Network neuroscience has constituted a more powerful methodology in precision medicine for DRE [14], [15], [16]. Causal brain network, defined as the directed influence (or causal driving effect) that one brain region or neuron exerts over another [17], has become a useful quantitative analysis in brain disease classification, pathological mechanism investigation, precise diagnosis and treatment, etc. [18], [19], [20], [21], [22], [23]. Seizure generation is believed to be a process driven by the epileptic network[24], [25], [26]; thus, understanding the causal connectivity within EZ, a clinically annotated region which are surgically resected, is crucial in revealing the triggering mechanism of epileptic seizures originating [27]. Characterizing causal connectivity in epileptic network would considerably facilitate precision medicine of DRE patients [15], [28], [29], [30], [31], [32].

### B. Problems Statement

The clinically annotated EZ is defined as the anatomical area to be treated (resection or ablation). In clinical neurosurgical procedures, DRE patients are subjected to long-term intracranial electroencephalography (iEEG) monitoring, typically such as electroencephalogram (ECoG), generally lasting for several days. The iEEG morphological and quantitative analysis combining with other clinical prior knowledge and multimodal clinical imaging data (i.e., MRI, PET) are collected comprehensively by several clinical experts to locate EZ for surgical targets. More than one year follow-up is conducted to determine surgical outcome, including successful surgery (seizure free with Engel I) and failed surgery (seizure recurring with Engel II∼IV). As shown in Fig. 1, a successful surgery removes the actual epileptic foci through the clinically annotated EZ. Whereas, a failed surgical treatment can not completely remove this epileptic focus, and the NEZ still contains epileptic foci. Meanwhile, the iEEG recordings in contact with the EZ do not visually differ significantly between successful and failed surgery. Therefore, quantitatively analyzing iEEG is clinically essential in localizing surgical targets and largely determining the DRE’s surgical outcomes.

**Fig. 1.**
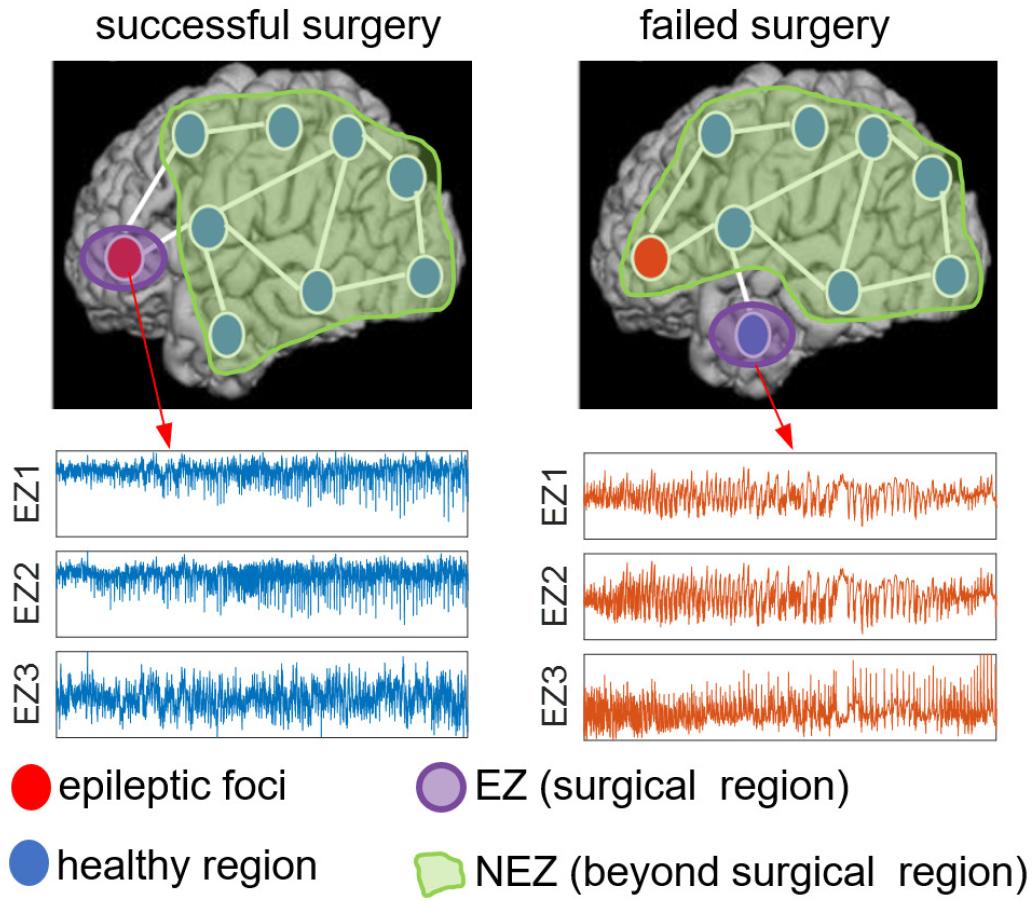
Example plots of successful and failed surgeries, and sample ictal ECoG signals of EZs.

As for the methodology, data-driven causal brain network algorithms mainly include Granger causality (GC), conditional Granger causality (cGC), kernel Granger causality (kGC), transfer entropy (TE), partial transfer entropy (pTE), symbolic transfer entropy (STE), convergent cross-mapping (CCM), etc. The advantages and disadvantages of the above methods have been discussed in detail in previous studies[23], [33], [34], [35], [36], [37], [38]. However, their performances in DRE analysis have not been explored in depth. A unified comparison on the same dataset is a necessary work to confirm which causal brain network achieves the best prediction of DRE surgical outcomes.

The brain network properties in EZ are significantly different from those in NEZ [9], meanwhile in different frequency bands of iEEG [15], which are worthy of grouping research. Some clinical covariates, such as DRE type (lesional and non-lesional patients), may potentially influence quantitative analysis conclusions regarding DRE. Unfortunately, systematic statistical analyses are still lacking. Some potential biomarkers for surgical outcomes should be discussed by grouping clinical covariates before their reliability can be confirmed.

### C. Contributions and Structure of This Study

Herein, we systematically explored ictal causal brain network closely related to surgical outcomes. Specifically, we included multicenter dataset, collected at two clinical centers and containing ECoG recordings from patients with lesional and non-lesional DRE. Based on the same dataset, six widely used data-driven algorithms were applied to construct the EZ and NEZ causal brain networks in multiple frequency bands, respectively. Mann-Whitney-U-test between successful and failed surgeries was performed and different causal algorithms were compared. In addition, clinical covariates were considered to facilitate the comprehensive statistical analysis and verify reliable biomarkers associated with surgical outcomes. Finally, based on the causal brain network features and machine learning algorithms, prediction models were established and verified by 5-fold cross validation. The developed prediction model was also compared with previous studies.

The remainder of this paper is structured as follows. Section II introduces the methods and materials. Section III explains statistical results of various causal brain networks. Section IV presents the prediction results of machine learning. Finally, Section VI discusses and concludes this study.

## II. Methods and materials

### A. Dataset Description

A public dataset [9], totally 17 DRE patients from two clinical centers with 55 seizures, was included in this retrospective cohort study. At all centers, ECoG data were recorded using Nihon Kohden or Natus acquisition system with a typical sampling rate of 1000Hz. For each DRE patient, the clinically annotated EZ is defined as the anatomical area to be treated (i.e., resected). Surgical outcomes were classified by epileptologists using the Engel Surgical Outcome Scale classification system. Successful surgical outcomes are defined as free of disabling seizures (Engel class E I) and failed outcomes as not free of disabling seizures (Engel classes, E II ∼ E IV) at 12+ months post-operation. Furthermore, the number of seizures per patient varies from 1 to 6. The detailed demographics and clinical information are listed in Tab. I.

**Tab. I.** Patient demographics and clinical information.

| Patient ID | Clinical center | Gender | Age (years) | MRI Pathology | Follow up/year | Engel class | Clinical complexity | # Seizure | Therapy | Implant type |
| --- | --- | --- | --- | --- | --- | --- | --- | --- | --- | --- |
| JH101 | JHU | / | / | Non-lesional | 1 | E IV | CC4 | 4 | Resection | ECOG |
| JH103 | JHU | / | / | Non-lesional | 1 | E IV | CC3 | 3 | Resection | ECOG |
| JH105 | JHU | / | / | Non-lesional | 1 | E I | CC2 | 4 | Resection | ECOG |
| JH108 | JHU | / | / | Non-lesional | 1 | E IV | CC4 | 6 | Resection | ECOG |
| Pt1 | NIH | F | 30 | Lesional | 3 | E I | CC1 | 4 | Resection | ECOG |
| Pt2 | NIH | F | 28 | Lesional | 3 | E I | CC1 | 3 | Resection | ECOG |
| Pt3 | NIH | M | 45 | Non-lesional | 2 | E I | CC3 | 2 | Resection | ECOG |
| Pt6 | NIH | M | 33 | Non-lesional | 3 | E II | CC4 | 1 | Resection | ECOG |
| Pt7 | NIH | F | 39 | Non-lesional | 7 | E III | CC3 | 3 | Resection | ECOG |
| Pt8 | NIH | M | 25 | Non-lesional | 2 | E I | CC3 | 3 | Resection | ECOG |
| Pt10 | NIH | F | 44 | Lesional | 1 | E II | CC1 | 3 | Resection | ECOG |
| Pt11 | NIH | M | 31 | Non-lesional | 1 | E I | CC3 | 4 | Resection | ECOG |
| Pt12 | NIH | F | 43 | Non-lesional | 5 | E II | CC4 | 2 | Resection | ECOG |
| Pt13 | NIH | M | 27 | Non-lesional | 2 | E I | CC3 | 4 | Resection | ECOG |
| Pt14 | NIH | F | 49 | Lesional | 1 | E IV | CC1 | 3 | Resection | ECOG |
| Pt15 | NIH | F | 59 | Lesional | 2 | E I | CC1 | 4 | Resection | ECOG |
| Pt16 | NIH | F | 52 | Lesional | 2 | E I | CC1 | 3 | Resection | ECOG |
| Pt17 | NIH | M | 13 | Lesional | 4 | E I | CC1 | 3 | Resection | ECOG |

Clinical center: JHU = Johns Hopkins University; NIH = National Institutes for Health Clinical Center. M = male, F = female;
Engel class E I = seizure free; E II = significant seizure frequency reduction; E III = slight seizure frequency reduction; E IV = no change after surgery.
Clinical complexity CC1 ~ CC4 is ordered by increasing localization difficulty: lesional (CC1), focal temporal (CC2), focal extratemporal (CC3) and multifocal (CC4). CC2~ CC4 are non-lesional.

**Tab. II.** Statistical results of all subjects in EZ regions.

| Frequency band | S Vs. F in EZ |  |  |  |  |  |  |  |  |  |  |  |
| --- | --- | --- | --- | --- | --- | --- | --- | --- | --- | --- | --- | --- |
|  | CCM |  | TE |  | pTE |  | sTE |  | GC |  | kGC |  |
| | $P$ value | Effect size | $P$ value | Effect size | $P$ value | Effect size | $P$ value | Effect size | $P$ value | Effect size | $P$ value | Effect size |
| Full | **5.00E-05** | **0.61** | 0.21708 | 0.19 | 0.07559 | 0.28 | 2.10E-04 | 0.56 | 0.05215 | 0.30 | 0.64393 | 0.07 |
| $\delta$ | **1.65E-03** | **0.48** | 0.19987 | 0.20 | 0.34751 | 0.15 | 4.16E-03 | 0.45 | 4.48E-03 | 0.44 | 0.07559 | 0.28 |
| $\theta$ | **7.85E-08** | **0.77** | 0.06100 | 0.29 | 0.11675 | 0.25 | 4.03E-03 | 0.44 | 0.05068 | 0.30 | 0.07442 | 0.28 |
| $\alpha$ | **1.75E-07** | **0.75** | 0.02169 | 0.36 | 0.85708 | 0.03 | 1.55E-03 | 0.48 | 0.01989 | 0.36 | 0.55669 | 0.09 |
| $\beta$ | **2.30E-04** | **0.56** | 0.14516 | 0.23 | 0.01742 | 0.37 | 0.04248 | 0.32 | 0.03572 | 0.33 | 0.38383 | 0.14 |
| $\gamma$ | **3.10E-04** | **0.56** | 9.00E-04 | 0.50 | 0.04521 | 0.31 | 0.01209 | 0.39 | 0.11675 | 0.25 | 0.50521 | 0.11 |
| HF | 1.01E-05 | 0.65 | **3.78E-07** | 0.74 | 0.08386 | 0.27 | 1.20E-04 | 0.58 | 0.03572 | 0.33 | 0.84480 | 0.03 |

**Tab. III.** Statistical results of all subjects in NEZ regions.

| Frequency band | S Vs. F in NEZ |  |  |  |  |  |  |  |  |  |  |  |
| --- | --- | --- | --- | --- | --- | --- | --- | --- | --- | --- | --- | --- |
|  | CCM |  | TE |  | pTE |  | sTE |  | GC |  | kGC |  |
| | $P$ value | Effect size | $P$ value | Effect size | $P$ value | Effect size | $P$ value | Effect size | $P$ value | Effect size | $P$ value | Effect size |
| Full | 0.62160 | 0.08 | **1.30E-03** | **0.49** | 1.65E-03 | 0.48 | 0.05262 | 0.30 | 0.22912 | 0.19 | 0.01522 | 0.38 |
| $\delta$ | 0.05668 | 0.30 | 0.41916 | 0.13 | 0.78839 | 0.04 | **1.66E-04** | **0.59** | 0.31114 | 0.16 | 0.78401 | 0.04 |
| $\theta$ | 0.34227 | 0.15 | 0.74813 | 0.05 | 0.72242 | 0.06 | **4.03E-03** | **0.44** | 0.27491 | 0.17 | 0.09284 | 0.26 |
| $\alpha$ | 0.95629 | 0.01 | 0.86939 | 0.03 | 0.41916 | 0.13 | 0.13241 | 0.24 | 0.26124 | 0.18 | 0.10598 | 0.25 |
| $\beta$ | 0.67802 | 0.07 | 0.91893 | 0.02 | 0.41016 | 0.13 | 0.09284 | 0.26 | 0.10948 | 0.25 | 0.35854 | 0.14 |
| $\gamma$ | 0.41916 | 0.13 | 0.06559 | 0.29 | 0.24804 | 0.18 | 0.63272 | 0.08 | **1.38E-03** | **0.49** | 0.04349 | 0.31 |
| HF | 9.54E-03 | 0.40 | 0.96877 | 0.01 | 0.53580 | 0.10 | 0.19987 | 0.20 | 0.18366 | 0.21 | **1.99E-05** | **0.63** |

### B. ECoG Preprocessing

Some study reported that quantified EZ using ictal markers were the only statistically significant determinants for surgical prognosis [39], therefore, for each ECoG recording, we uniformly intercepted the 10-s ictal signal. Before estimating the causal brain network, the intercepted ECoG was bandpass filtered between 0.5 and 300 Hz with a fourth-order Butterworth filter, and notch filtered at 60 Hz and its harmonics with a stopband of 2 Hz. In addition, ECoG channels deemed ‘bad contacts’ (e.g., broken or excessively noisy or artifactual) by the clinicians were discarded. Then ECoG data was filtered as six different frequency bands: δ (0.4 ∼ 4 Hz), θ (4 ∼ 8 Hz), α (8 ∼ 13 Hz), β (13 ∼ 30 Hz), γ (30 ∼ 80 Hz), high frequency oscillation (HF, >80Hz). Standardized z-scores were also used to normalize the variance of the ECoG data in each channel. Finally, we computed the ictal causal brain network based on the preprocessed multiband ECoG within EZ and NEZ, obtaining the δ-, θ-, α-, β-, γ-, HF-EZ networks and NEZ networks. In addition, the preprocessed unfiltered full-band ECoG were also estimated to obtain the brain networks.

### C. Algorithms for Measuring Causal Brain Network

In this paper, we used six data-driven causal brain network algorithms for comparative study, including GC, kGC, TE, pTE, sTE and CCM. GC is a statistical hypothesis test used to determine whether one time series can be used to predict another, which definition is based on the variance of the best least squares prediction using all the information at some point in the past. In 2008, D.Marinazzo et al. generalize GC to the nonlinear case using the theory of reproducing kernel Hilbert spaces, named kGC, of which the nonlinearity of the regression model can be controlled by choosing the kernel function. TE is used in information theory and time series analysis to quantify the directional information flow between two stochastic processes, which can help predict the future states of another process from one process. Unlike GC, TE can capture non-linear dependencies without a linear relationship assumption. pTE is an extension of transfer entropy, which is proposed to detect causal effects among observed interacting systems, and particularly to distinguish direct from indirect causal effects. In sTE, Staniek et al. defined symbols by reordering the amplitude values of time series to estimate TE. They demonstrated sTE is a robust and computationally fast method to quantify the dominating direction of information flow between time series. CCM is a nonlinear method used to assess causality and reveal directional relationships between variables in a dynamical system based on time series data. CCM develops in the context of complex systems analysis and aims to reveal causal influences between variables even in the absence of a clear mechanistic understanding.

## III. Statistical results of causal brain network

### A. Statistical results of various causal brain networks between EZ and NEZ

In this study, we calculated causal brain networks for all 59 seizures with different causal coupling algorithms (CCM, TE, pTE, sTE, GC, and kGC), including the causal coupling strength between iEEG signals in all frequency bands within the EZ regions and the NEZ regions.

We grouped the results of EZ and NEZ regions into successful surgical group and failed surgical group for statistical analysis, respectively. Mann-Whitney U test was used for statistical analysis, and P value and Cohen’s d size were used for evaluation. The statistical results in Fig. 2 and 3 showed that the causal coupling strength in EZ regions calculated by CCM and sTE methods of the successful surgical group was significantly higher than that of failed surgical group at all frequency bands. Moreover, the results of CCM method have higher significance compared to sTE, especially in the θ band (smallest *p* value: 7.85e-8, largest effect size: 0.77). The results of GC, TE and pTE methods have significant differences in 4 frequency bands (δ-, α-, β-, γ-iEEG network), 3 (α-, γ-, HF-iEEG network) and 2 frequency bands (β-, γ-iEEG network) respectively, while that of kGC showed no significance in all frequency bands. In NEZ regions, there are fewer significantly different results than that in EZ. In α- and β-bands, the results of all methods do not have significant differences. In other frequency bands, some methods have significant results, but the performance is not as good as EZ region.

**Fig. 2.**
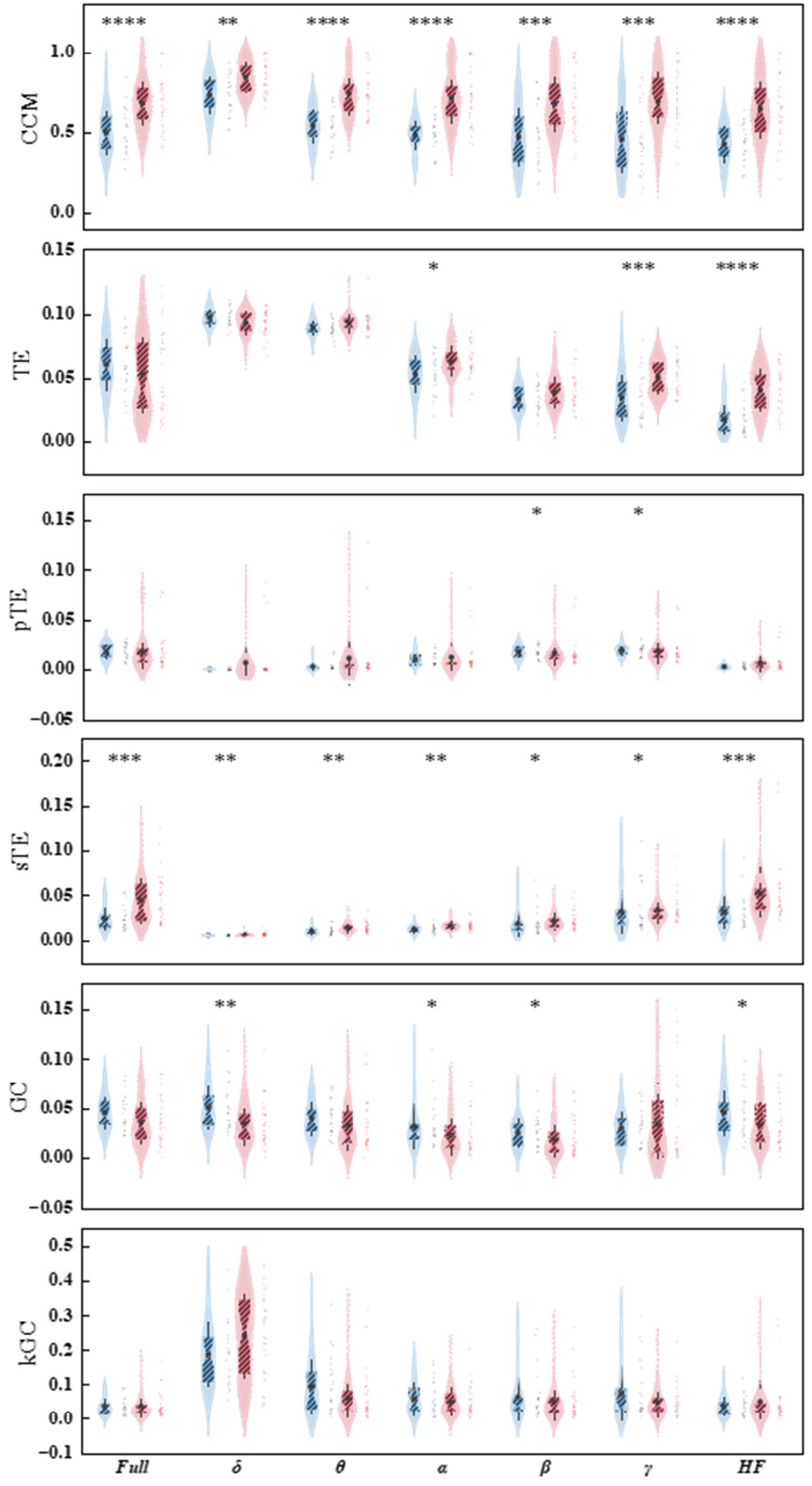
Statistical results among various methods in EZ regions. The blue violin plots represent failed group and the red represent the successful group. The marker “^*^” indicates the P value through Mann-Whitney-U-test. *: *P* < 0.05; ^**^: *P* < 0.01; ^***^: *P* < 0.001; ^****^: *P* < 0.0001.

**Fig. 3.**
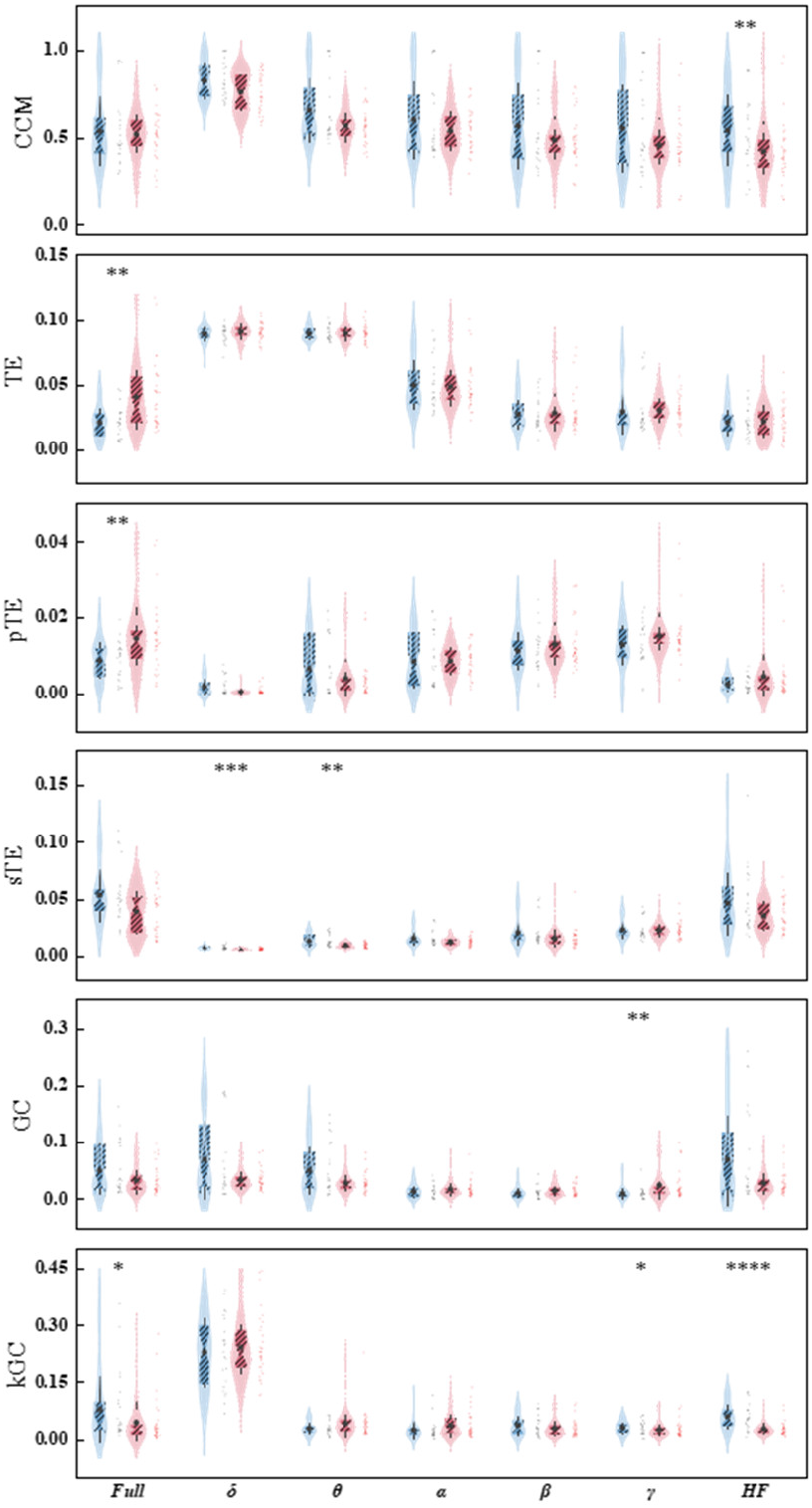
Statistical results among various methods in NEZ regions. The blue violin plots represent failed group and the red represent the successful group. The marker “^*^” indicates the *P* value through Mann-Whitney-U-test. ^*^: *P* < 0.05; ^**^: *P* < 0.01; ^***^: *P* < 0.001; ^****^: *P* < 0.0001.

### B. Statistical results considering different clinical covariates

In order to make the results more reliable and avoid as much bias as possible due to a single data source, we combined two data sets from different clinical centers in this study. And in this combined dataset, there are two different types of DRE patients. Therefore, we conducted statistical analysis on samples from different clinical centers and different types of DRE patients.

First, we performed a statistical analysis of the outcomes of the successful and failed surgical groups in the two datasets. The results are shown in Fig. 4. In the JHU dataset, the causal coupling strength of the EZ region calculated by GC was significantly higher in the successful group than that in the failed group at all frequency bands. In the NIH dataset, the CCM method showed the best performance. At the same time, in the NIH dataset, the causal coupling strength of GC methods in multiple frequency bands (except α-band) in the NEZ region also showed significant differences between the two groups. In addition, we found that the HF-band had the most significant results in both the JHU and NIH datasets.

**Fig. 4.**
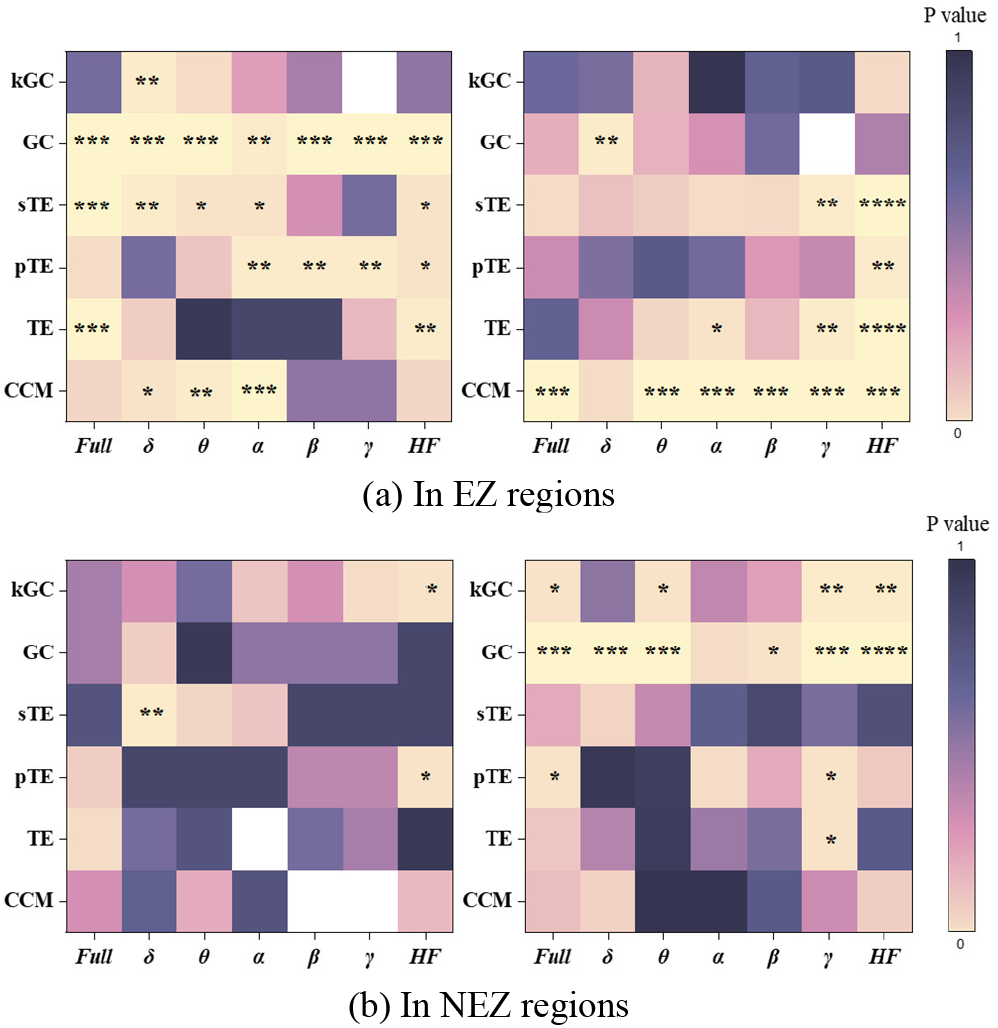
P value heatmap of statistical results among various methods (a) in EZ regions and (b)in NEZ regions from two clinical centers: JHU and NIH. The left plots show the results of JHU and the right show the results of NIH. The marker “*” indicates the P value through Mann-Whitney-U-test. ^*^: *P* < 0.05; ^**^: *P* < 0.01; ^***^: *P* < 0.001; ^****^: *P* < 0.0001.

Finally, we performed a statistical analysis in different types of DRE patients. In the subdataset composed of non-lesional DRE (NL-DRE), the causal coupling strength of the successful surgical group calculated by CCM and sTE in EZ regions were significantly higher in all frequency bands than that of the failed surgical group. While in NEZ regions, there were less significances. In the lesional DRE (L-DRE) subdataset, CCM also showed a good distinction effect. In EZ regions, CCM results showed significant differences between successful and failed groups in most frequency bands (except δ-band). However, the number of significant results in NEZ regions was similar to those in EZ regions in the L-DRE subdataset. The results are shown in Fig. 5.

**Fig. 5.**
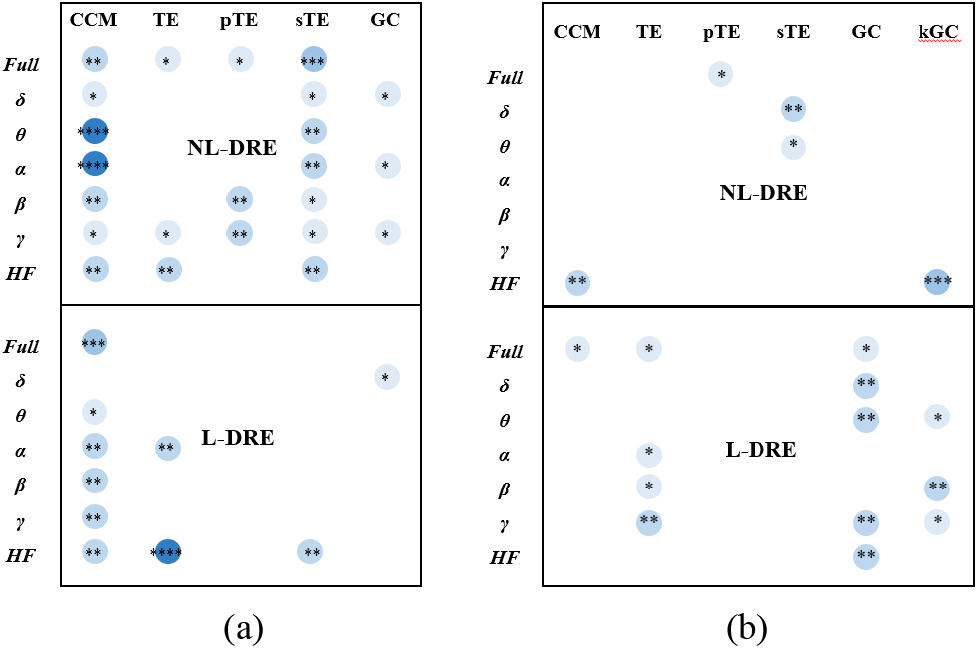
Statistical results of NL-DRE and L-DRE sub dataset. (a)In EZ regions, (b)In NEZ regions.

## IV. Surgical outcome prediction results based on machine learning

For the surgery outcome prediction, we undertook the classification between surgery success and failure group. Based on the statical analysis, we extracted these brain network features showing significant differences between successful and failed surgeries, calculated by CCM, TE, pTE, sTE, GC, kGC. EZ and NEZ brain network features were considered separately. Four classifiers, support vector machine (SVM), Naive Bayes (NB), logistic regression (LR) and linear discriminant analysis (LDA) were applied in the prediction. The kernel function for the SVM was specified as a Gaussian kernel and we employed Bayesian optimization to modify the SVM’s and NB’s hyperparameters during cross-validation. To assess the predicting performances, 5-fold cross-validation was employed. The average accuracy results are presented in Tab. V. The SVM model with Gaussian kernel function and Bayesian Optimization exhibits the best performance, achieving an average accuracy of 84.48%. Although kGC-related NEZ brain network features also achieve good performance, the best average accuracy is still below 80% (79.31% in SVM classifier).

Moreover, our work was compared with previous studies in predicting DRE’s surgical outcomes. The performance comparisons are listed in Tab. VI. In general, the EZ brain network features calculated by CCM combined with SVM classifier achieved the best overall performances, i.e., accuracy = 84.48%, PPV = 75.86%, NPV = 93.10%, sensitivity = 91.67%, specificity = 89.92% and F1-score = 83.02%.

**Tab. IV.** Average accuracy using 5-fold cross-validation.

| Classifier | EZ brain network features including frequency bands with significant difference |  |  |  |  |  |
| --- | --- | --- | --- | --- | --- | --- |
| | CCM<br>(Full, $\delta$, $\theta$, $\alpha$, $\beta$, $\gamma$, HF) | TE<br>($\alpha$, $\gamma$, HF) | pTE<br>($\beta$, $\gamma$) | sTE<br>(Full, $\delta$, $\theta$, $\alpha$, $\beta$, $\gamma$, HF) | GC<br>($\delta$, $\alpha$, $\beta$, HF) | kGC |
| SVM | **84.48%** | 70.69% | 58.62% | 79.31% | 75.86% | \ |
| NB | **77.59%** | 63.79% | 67.24% | 68.97% | 74.14% | \ |
| LR | **79.31%** | 67.24% | 58.62% | 72.41% | 60.34% | \ |
| LDA | **77.59%** | 70.69% | 58.62% | 75.86% | 60.34% | \ |
| Classifier | NEZ brain network features including frequency bands with significant difference |  |  |  |  |  |
| | CCM<br>(HF) | TE<br>(Full) | pTE<br>(Full) | sTE<br>($\delta$, $\theta$) | GC<br>($\gamma$) | kGC<br>(Full, $\gamma$, HF) |
| SVM | 68.97% | 70.69% | 72.41% | 70.69% | 68.97% | **79.31%** |
| NB | 70.69% | 65.52% | 70.69% | **62.07%** | 60.34% | **77.59%** |
| LR | 65.52% | 62.07% | 62.07% | 62.07% | 62.07% | **77.59%** |
| LDA | 67.24% | 60.34% | 62.07% | 63.79% | 60.34% | **77.59%** |

**Tab. V.** The comparison of prediction performance between this paper and previous studies.

| Ref. | Data | MRI | Feature/Methodology | Accuracy | PPV | NPV | Sensitivity | Specificity | F1-score |
| --- | --- | --- | --- | --- | --- | --- | --- | --- | --- |
| **This paper** | **EEG** | **N/P** | **causal strength in EZ** | **84.48%** | **75.86%** | **93.10%** | **91.67%** | **89.92%** | **83.02%** |
| [9] | iEEG | N/P | Neural fragility, structured RF | 76% | 90.3% | 87.2% | \ | \ | \ |
| [13] | iEEG | N/P | functional connectivity | 45~72% | 66~100% | 31~66% | \ | \ | \ |
| [40] | iEEG | N/P | spike propagation in source space | 69% | 79% | 56% | \ | \ | \ |
| [41] | EEG/MEG | N/P | functional connectivity | 66~76% | 65~81% | 63~76% | \ | \ | \ |
| [42] | PET/MRI | N | feature of hypometabolism | 70~80% | 83~87% | 53~72% | 69~86% | 61~73% | \ |
| [43] | PET/FDG | N | individual metabolic connectome | 83.59% | \ | 75.00% | 92.79% | \ | \ |
| [44] | SEEG | P | high-frequency oscillations | 69.5% | 60.7% | 74.7% | 58.6% | 76.3% | \ |
| [45] | EEG/MRI | N/P | ripple onset | 71% | 91% | 59% | \ | \ | \ |
| [46] | EEG/MRI | N/P | Logistic regression | 72% | \ | \ | \ | \ | \ |
| [47] | foramen ovale EEG | N/P | power spectral densities/Logistic Regression | 68% | \ | 57% | \ | \ | \ |
| [48] | SEEG | P/N | SEEG desynchronization | \ | \ | \ | 89.3% | 82.4% | \ |

## V. Discussion and conclusion

### A. Implications for epileptic causal brain network

Brain network methodology has effectively promoted the study of drug-resistant epilepsy (DRE). Some connectome biomarkers have been gradually developed to evaluate DRE’s surgical outcomes [13], [29], [40], [41], [43], [49], [50], [51], [52]. Causal brain network can reveal the triggering mechanism of epileptic seizures originating [27], and has considerably facilitate DRE studies [15], [28], [29], [30], [31], [32].

In this study, a multi-center ECoG dataset was included, and causal brain network closely related to surgical outcomes was systematically explored. A series of comparative results together support our conclusion: CCM-defined EZ brain network is the most effective biomarker for predicting DRE’s surgical outcomes. For successful surgeries, causal connectivity in EZ is significantly higher than that of failed surgeries. The findings are consistent with previous study [27], seizure-onset regions demonstrate high inward directed connectivity. One possible explanation is that the EZ forms the driving source of an unusually complex system, whose interior is strongly coupled, facilitating seizures and propagation. For the failed surgeries, their EZ (clinically annotated regions to be treated), are not true surgical foci partly or totally, so their causal connectivity is unusually weak compared to those of true epileptogenic foci. The NEZ brain network was less effective in distinguishing surgical outcomes, only showing limited significant differences, while the *P*-value was greater than that of EZ brain network. Except for CCM, the kGC algorithm represented the best performance, because kGC is based on kernel methods with good quantification accuracy and robustness, and it can effectively quantify nonlinear causal coupling, which is an important feature in complex brain network [53], [54]. Similarly, in machine learning prediction, kGC-feature based models also exhibit suboptimal performance of 79.31% accuracy. Considering clinical covariate of DRE types, i.e., lesional and non-lesional DRE, causal brain network results were statistically analyzed separately. CCM-defined EZ causal connectivity showed more significant differences between successful and failed surgeries of non-lesional DRE patients, with *P* <0.0001in θ- and α-band network, while, *P* < 0.001 in Full-band network in distinguishing lesional DRE’s surgical outcomes. The lesional DRE’s brain network generate a heterogeneous complex system. When CCM quantifies the causal coupling in complex system, it is based on the non-separability of system variables, and more attention is paid to the system’s homogeneity.

### B. Limits and future studies

It should be pointed out that there are still several limitations in current study. The surgical outcomes of neurosurgery are only divided to two types, i.e., success (Engel I) and failure (Engel II∼IV). For further research, a possible improvement could involve transforming the binary classification into a four-class classification problem to achieve a more nuanced categorization. What’s more, the statistical results of lesional and non-lesional patients demonstrate that there exist some differences in different types of patients. As a result, subsequent steps could involve employing transfer learning methods to achieve model generalization across different types of patients, or constructing a multi-task model, initially classifying and predicting the type of the patients inputted and then predicting the surgical outcomes. The dataset of this study remains relatively limited. Future work will recruit more DRE subjects and may extend to multicenter clinical studies, including more types of DRE (CC1 ∼ CC4), and more iEEG recordings, i.e., ECoG and stereoelectroencephalogram (SEEG).

### C. Conclusion

Causal brain network in patients with drug-resistant epilepsy was systematically investigated to evaluates surgical outcomes in this study. Multicenter ECoG datasets containing 17 DRE patients with 55 seizures were included, and EZ- and NEZ-causal connectivity involving multi frequency bands (δ, θ, α, β, γ, HF, Full) was calculated by 6 different algorithms. Mann-Whitney-U-test show that α-EZ brain network calculated by CCM gains the most significant differences between successful and failed surgeries, with a *P* value of 7.85e-08. Combining brain network features and machine learning, prediction models were established, and a SVM classifier demonstrates the highest average accuracy of 84.48% through 5-fold cross validation. The findings indicates that the CCM-defined EZ brain network is a reliable biomarker for predicting DRE’s surgical outcomes.

